# The Colonic Mucus Layer is Thinner and is Associated with Goblet Cell Hyperplasia in the *db/db* Mouse Model of Type 2 Diabetes

**DOI:** 10.64898/2026.04.02.716104

**Authors:** Matthew C. Rowe, Matthias Demuynck, Abhipree Sharma, Cameron J. Nowell, Caitlin Owyong, Nimna Perera, N. Jennifer Tang, Nicholas A. Veldhuis, Pradeep Rajasekhar, Rebecca H. Ritchie, Miles J. De Blasio, Simona. E. Carbone, Daniel P. Poole

## Abstract

**Background & Aims:** Diabetes mellitus has been associated with both intestinal barrier dysfunction and peripheral neuropathy leading to increased risk of infection. The mucus layer forms a physical barrier against pathogens and is a critical component of the intestinal barrier that may be impaired in diabetes. This study aimed to assess how diabetes impacts goblet cells (GCs), mucus layer integrity, and innervation in the colon.

**Methods:** Fluorescence microscopy was used to investigate GCs, the mucus layer, and innervation in the colon of *db/db* mice. Custom open-access image analysis pipelines were developed to quantify GC numbers, location and content, mucus thickness, bacterial colonization, and innervation density in intestinal tissue sections. We also treated mice with the clinically used glucagon-like peptide 1 receptor (GLP-1R) agonist liraglutide to assess its capacity to reverse pathological changes to GCs and the mucus layer in a model of established type 2 diabetes (T2DM).

**Results:** The mucus layer was significantly thinner in the colon of *db/db* mice with established diabetes and bacteria more readily colonized the epithelium and crypts. Intercrypt GC numbers were significantly reduced in *db/db* mice. However, there were significantly more GCs per crypt, and crypts were elongated in the *db/db* colon. Innervation was reduced in the mucosa and external muscle of the colon, consistent with diabetic neuropathic changes. Liraglutide treatment increased the size of GCs but had no effect on GC numbers, mucus thickness, or innervation in this model of established T2DM.

**Conclusions:** Mucus barrier dysfunction and GC hyperplasia is evident in the *db/db* colon. Increased microbial penetrability through the mucus layer suggests potential implications for the increased risk of gastrointestinal infection in diabetes.

## Introduction

Diabetes mellitus (DM) is a metabolic disease caused by defects in insulin secretion or function. DM is defined by persistent hyperglycemia caused by defects in insulin secretion or function. There are two major types of DM. Type 1 diabetes (T1DM) is an autoimmune condition that results from destruction of pancreatic β cells by the immune system, stopping the production of insulin. Type 2 diabetes (T2DM) is caused by insulin resistance or insufficient insulin production. Beyond the dysregulation of glycemic control, DM is associated with significant gastrointestinal and neural complications including gastroparesis and sensory and enteric neuropathy^1,2^. Gastrointestinal dysfunction is best studied in the context of gastroparesis, and is characterized by delayed gastric emptying and mechanical obstruction of the stomach due in part to autonomic denervation^3^. The nervous system also has a critical role in regulating broader gastrointestinal functions including maintenance of intestinal barrier integrity. Therefore, intestinal barrier functions are likely to be impacted in DM. The intestinal barrier is comprised of a monolayer of epithelial cells, tight junctions, and a mucus layer that selectively protects the epithelium from pathogens and physical damage. DM is often associated with increased intestinal permeability, thought to be due to a disrupted intestinal barrier^4–6^. This aligns with the increased susceptibility of patients with DM to infection and chronic intestinal inflammation^7,8^. However, the extent to which the intestinal barrier is impaired in DM, particularly the mucus layer, is poorly understood.

An effective intestinal barrier is critical for maintaining intestinal homeostasis. Increased intestinal permeability in DM is best evidenced by changes in tight junction proteins. Altered zonulin expression in serum has been correlated with increased intestinal permeability to macromolecules in DM patients^9–11^. Additionally, occludin transcription was also elevated in T1DM duodenal patient biopsies^10^. However, a study investigating transcriptional changes to tight junction markers in the human T1DM small intestine reported no difference in occludin, zonulin or claudin expression^9^. In a high fat diet with streptozotocin model of T2DM, both occludin and zonulin expression were reduced in the duodenum, emphasizing the diversity in reported outcomes^12^.

The mucus layer is a physical barrier that prevents the translocation of bacteria to the epithelium^13^. Intestinal mucus is synthesized and secreted by goblet cells (GCs) in response to physical, chemical, and biological stimuli, including many neurotransmitters^14,15^. Disruption to GC functions may impair mucus layer homeostasis, leaving the epithelium more vulnerable to bacterial translocation^16^. Alterations to GCs and mucus secretion have been documented in DM. Mucus secretion induced by cholinergic stimulation was significantly reduced in the streptozotocin model of T1DM rat intestine, suggesting a deficit in GC function^17^. Additionally, transcription of mucus-associated genes including mucin-2 (MUC2) is reduced in the small intestine of T1DM patients and mouse models^10,18^. T1DM-associated effects on intestinal GC numbers appear to be highly variable, with reports that Periodic Acid-Schiff+ GCs are increased, decreased or unchanged^19–21^. The diversity of findings and limited equivalent investigation in T2DM indicate the need for further investigation. Understanding how the mucus layer and GCs are altered in DM may help to unravel the extent of intestinal barrier dysfunction that occurs and provide a possible explanation for the greater susceptibility to infection.

Neural signaling is an important regulator of GC function and mucus secretion. The intestine is innervated by the intrinsic enteric nervous system and also receives extensive extrinsic input from sensory and autonomic nerves, all of which can be disrupted in diabetic neuropathy^2,22^. Diabetic neuropathy is thought to be caused by prolonged hyperglycemia and impacts approximately 66% of DM patients^23^. It is commonly associated with symptoms of peripheral numbness and pain at the extremities^22,23^. Additionally, diabetic neuropathy has been correlated with elevated intestinal permeability and bacterial overgrowth compared to T1DM patients without autonomic neuropathy, suggesting that neuronal impairment may exacerbate intestinal barrier dysfunction^24,25^. This observation supports the concept that impaired neural signaling may be a key contributor to intestinal barrier dysfunction in DM patients.

Glucagon-like peptide 1 receptor agonists (GLP-1RA) are one of the common therapeutics prescribed to T2DM patients to regulate blood glucose levels. Activation of GLP-1R on pancreatic β cells stimulates endogenous insulin secretion to restore glycemic control^26^. GLP-1R is also widely expressed throughout the body, including in the gastrointestinal tract, and endogenous GLP-1 is primarily secreted by intestinal L-cells^27–29^. Beyond glycemic regulation, GLP-1 has an important role in regulating gastrointestinal functions including nutrient absorption, energy metabolism, and gastric motility^30^. Additionally, GLP-1RAs have neuroprotective effects in diabetes models and on cultured enteric neurons. More recently, GLP-1 has been implicated in the homeostatic regulation of the intestinal barrier. Liraglutide, a clinically used GLP-1RA significantly attenuates colonic permeability following acute LPS challenge in rats^31^. A single dose of liraglutide has also been shown to increase mucin transcription in the mouse duodenum^32^. Additionally, GLP-1R knockout was associated with reduced GC numbers in the colon of germ-free mice assessed by MUC2 immunolabeling^33^. Prior studies support a role for GLP-1R activation in maintaining intestinal barrier integrity and homeostatic regulation of GC function.

In this study, we investigate the impact of diabetes on colonic GCs, the mucus layer and innervation using the *db/db* model of T2DM. We demonstrate that the number of intercrypt GCs are reduced, whereas the total number of GCs per crypt is increased in the *db/db* colon. Furthermore, the mucus layer is significantly thinner, and bacteria more readily colonize the colonic epithelium and crypts in T2DM, consistent with a defective mucus barrier. Innervation is also notably reduced throughout the colon. Additionally, we assessed the potential for liraglutide to maintain the mucus layer and GCs in an established model of T2DM. GLP-1RA treatment was ineffective at reverting T2DM-associated changes to GC numbers, mucus thickness or innervation back to the levels of control mice, although it increased the average size of GCs.

## Results

### Goblet cells numbers and size are unchanged in the *db/db* colon

The mucus layer is an essential physical barrier that protects the epithelium. The effectiveness of this barrier is likely to be compromised in T2DM. T2DM-associated effects on colonic GCs and the mucus layer were assessed in wild type (WT), *db/h* (heterozygote littermate control), and *db/db* mice. Colons were fluorescently labeled with UEA1 and WGA to identify GCs and mucus **(Fig. 1A)**. UEA1 binds to fucose-residues detected in GCs towards the upper crypt, while WGA labels N-acetyl-D-glucosamine and sialic acid residues detected in GCs at the base of the crypt^34^. GCs were categorized into three groups: UEA1^+^/WGA^-^ (UEA1^+^ GCs), UEA1^-^/WGA^+^ (WGA^+^ GCs), and UEA1^+^ and/or WGA^+^ (Total GCs) **(Fig. S2)**. The total number of colonic GCs was unchanged in *db/db* mice compared to WT and *db/h* controls **(Fig. 1B)**. Similarly, there was no difference in the number of UEA1^+^ and WGA^+^ GCs in *db/db* mice compared to controls **(Fig. 1C, D)**. Intercrypt GCs reside at the luminal surface and rapidly secrete mucus under homeostatic conditions to maintain the mucus layer^34,35^. The number of UEA1^+^ intercrypt GCs was significantly decreased in *db/db* mice **(Fig. 1E)**. Treatment of diabetic *db/db* mice with liraglutide resulted in reductions in both the total number of GCs and UEA1^+^ GCs **(Fig. 1B, C)**. In contrast, there was no effect on the number of WGA^+^ or UEA1^+^ intercrypt GCs **(Fig 1D, E)**. Although GC size was unchanged in *db/db* mice, treatment of these mice with liraglutide was associated with a significant increase in the average size of all GCs **(Fig. 1F)**, UEA1^+^ GCs **(Fig. 1G)** and WGA^+^ GCs **(Fig. 1H)**. Positively labeled area of the major intestinal mucin, MUC2, in *db/db* mice was the same as controls **(Fig. 1I)**. Collectively, these findings indicate that there is a significant decrease in the number of intercrypt GCs in the colon of *db/db* mice, although the total number and size of GCs is unchanged.

**Figure 1:**
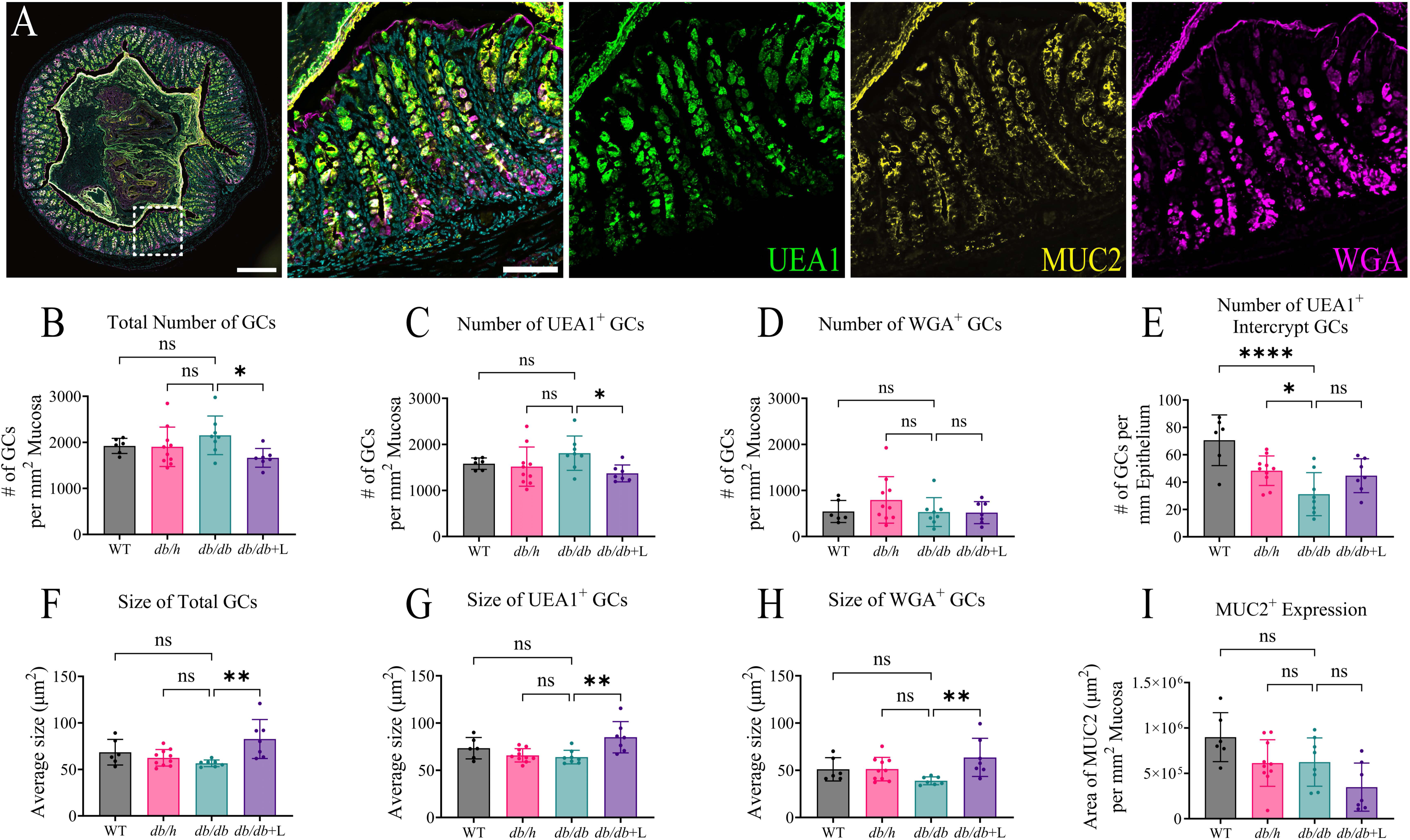
Colonic goblet cell properties are unaltered in db/db mice. (A) Representative image of a *db/h* mouse colon cryosection labelled for DAPI (cyan), UEA1 (green), MUC2 (yellow) and WGA (magenta). Diabetes had no effect on the number of (B) Total, (C) UEA1^+^, (D) or WGA^+^ GCs in WT, *db/h*, *db/db*, and *db/db*+liraglutide-treated mice. (E) The number of UEA1^+^ intercrypt GCs was reduced in the diabetic colon. Diabetes had no effect on the average size of (F) total, (G) UEA1^+^, and (H) WGA^+^ GCs. The average size of GCs was increased in liraglutide-treated *db/db* mice. (I) The area of MUC2^+^ expression within the mucosa was unchanged. Data are presented as the mean ± SD, with each data point representing the mean of triplicate measurements from an individual mouse. Data were analyzed using a one-way ANOVA with Dunnett’s multiple comparisons test. Scale bar: (A) 300µm, 75µm inset. ns=not significant; n=6-10 mice per group. *p<0.05; **p<0.01; ****p<0.0001.

### Goblet cell numbers are increased in the colonic crypts of *db/db* mice

GCs and other crypt-specific cell types may not be accurately represented in image analysis due to the inclusion of incomplete crypt structures and lamina propria tissue due to tissue sectioning artifacts. To exclude partial crypts and the lamina propria in GC measurements, analysis was restricted to full-length intact crypts from each animal group **(Fig. 2A, S2, S4)**. In contrast to analysis of the whole mucosa outlined in **Fig. 1**, the total number of GCs and the number of UEA1^+^ GCs per crypt were significantly increased in *db/db* tissues compared to WT and *db/h* controls **(Fig. 2B, C)**. Liraglutide treatment reduced the total number of GCs and the number of UEA1^+^ GCs per crypt back to control numbers at the 90% confidence level. The number of WGA^+^ GCs was unchanged **(Fig. 2D)**. Liraglutide treatment significantly increased the average GC size in the total, UEA1^+^ and WGA^+^ GC populations compared to the *db/db* group **(Fig. 2E, F, G)**. There was no difference in the average size of GCs between *db/db* and *db/h* or WT controls. The average length of crypts was increased in the *db/db* colon relative to *db/h* and WT controls **(Fig. 2H)**. Liraglutide treatment had no effect on this increase. UEA1^+^ GCs were distributed throughout the crypts with the mean location near the middle of the crypt **(Fig. 2I, N)**. The average location of UEA1^+^ GCs was shifted significantly towards the base of the crypts in *db/db* mice. WGA^+^ GCs were distributed towards the base of the crypt **(Fig. 2J, O)**. The mean location of WGA^+^ GCs was shifted significantly further towards the base of the crypts in *db/db* mice. There was no change to total area of MUC2^+^ labeling within entire crypts or specifically within GCs in the *db/db* group compared to WT or *db/h* controls **(Fig. 2K, L)**. Liraglutide treatment reduced the area of total MUC2^+^ labeling and MUC2^+^ labeling specifically within GCs at the 90% confidence level (p=0.0693). There was no difference to the total number of GCs when normalized to a standardized crypt length **(Fig. 2M)**. In summary, these data demonstrate that the increased number of GCs per crypt is correlated with elongated crypts in the diabetic colon.

**Figure 2:**
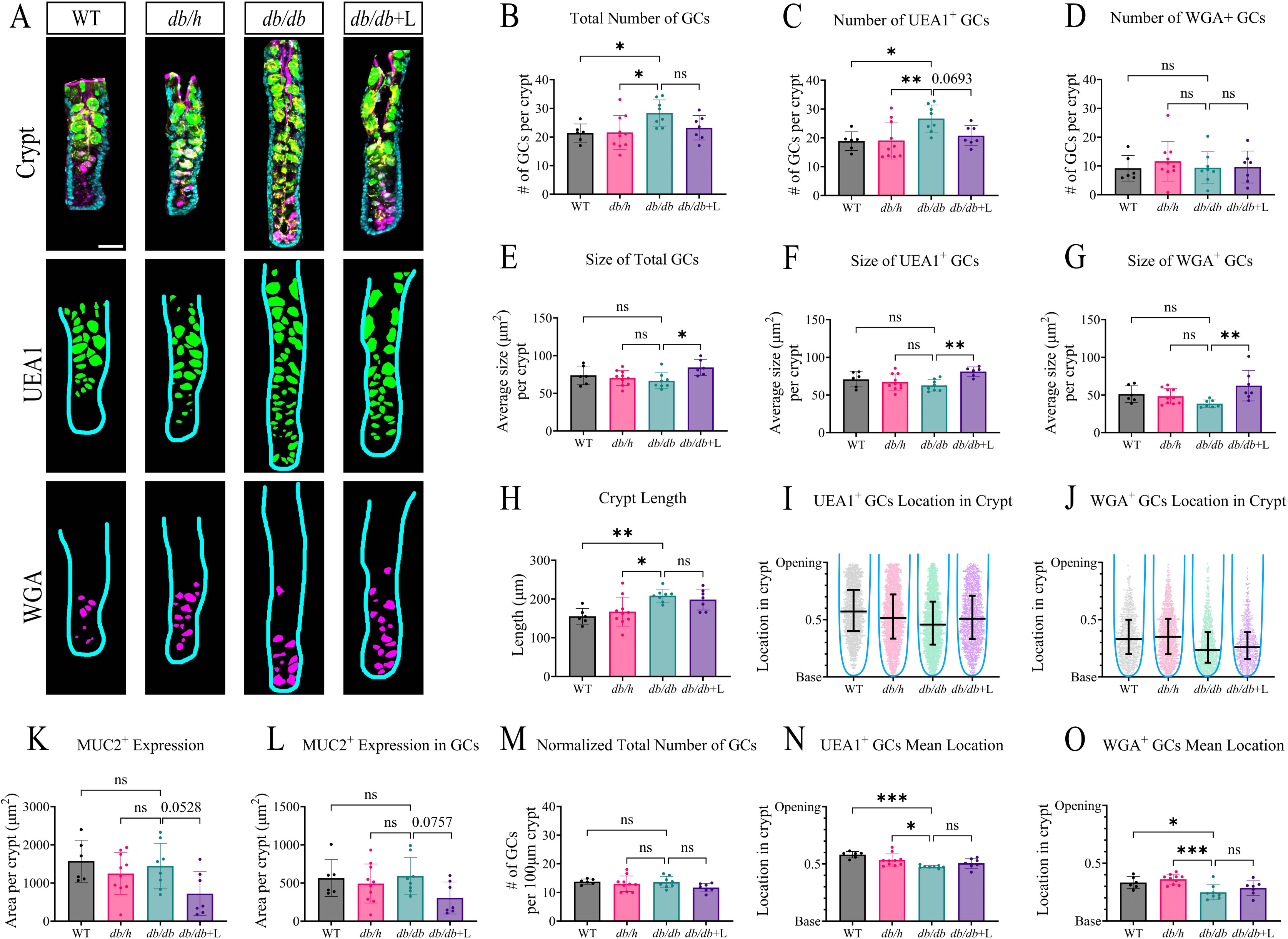
The number of goblet cells increases in the colonic crypts of db/db mice. (A) Representative images of colonic crypts labelled with DAPI (cyan), UEA1 (green), MUC2 (yellow) and WGA (magenta). Each crypt was digitally cropped to display segmented UEA1 (green) and WGA (magenta) positive GCs within an individual colonic crypt of WT, *db/h*, *db/db*, and *db/db*+liraglutide-treated mice. Diabetes increased the number of (B) Total and (C) UEA1^+^ GCs per crypt. (D) Diabetes had no effect on the number of WGA^+^ GCs per crypt. There was no difference in the average size of (E) Total (F) UEA1^+^, and (G) WGA^+^ GCs per crypt in the *db/db* colon compared to *db/h* and WT controls. Diabetic mice treated with liraglutide had larger GCs per crypt. (H) The average length of intact full-length crypts was significantly increased in the *db/db* colon. Mean distribution of (I) UEA1^+^ and (J) WGA^+^ GCs between the base and opening of crypts. Each data point represents an individual GC measured within a crypt. (K) Area of MUC2^+^ expression per crypt. (L) Area of MUC2^+^ expression within UEA1^+^ and/or WGA^+^ GCs per crypt. (M) The total number of GCs normalized per 100µm crypt length was unchanged in the db/db colon. Average distance of (N) UEA1^+^ and (O) WGA^+^ GCs from the base of the crypt. Data are presented the mean ± SD, with each data point represented the mean of triplicate measurements from an individual mouse. Data were analyzed using a one-way ANOVA with Dunnett’s multiple comparisons test. Scale bar: (A) 30µm. ns=not significant; n=6-10 mice per group. *p<0.05; **p<0.01; ***p<0.001.

### The colonic mucus layer is thinner in *db/db* mice and is associated with increased bacterial colonization

Mucus secretion by GCs is disrupted in the streptozotocin rat model of T1DM^17^. Additionally, bacteria colonization is reportedly increased in the small intestine of DM patients^36^. The colonic mucus layer was significantly thinner in the *db/db* colon compared to WT and *db/h* controls **(Fig. 3A, B)**. Liraglutide treatment had no effect on mucus layer thickness compared to untreated *db/db* mice. The distribution of DAPI^+^ bacteria-like cells in crypts and at the epithelial surface was assessed as a surrogate measure to mucus barrier function. In the *db/h* colon bacteria were present in the mucus layer and but were rarely observed at the epithelium or in the crypts. In marked contrast, there was extensive bacterial localization at the epithelial surface and DAPI^+^ biofilm formations within the crypts in the *db/db* colon **(Fig. 3C, D, E)**. Liraglutide treatment did not prevent the localization of bacteria at the epithelium or within the crypts. Increased bacterial association with the epithelium has been correlated with colitis and suggests disrupted mucus function^37,38^. Furthermore, a thinner colonic mucus layer has also been associated with ulcerative colitis in patients^39,40^. Together, our findings suggest that there is a diminished mucus layer and increased bacterial localization at the epithelium in T2DM, consistent with impaired mucus barrier function.

**Figure 3:**
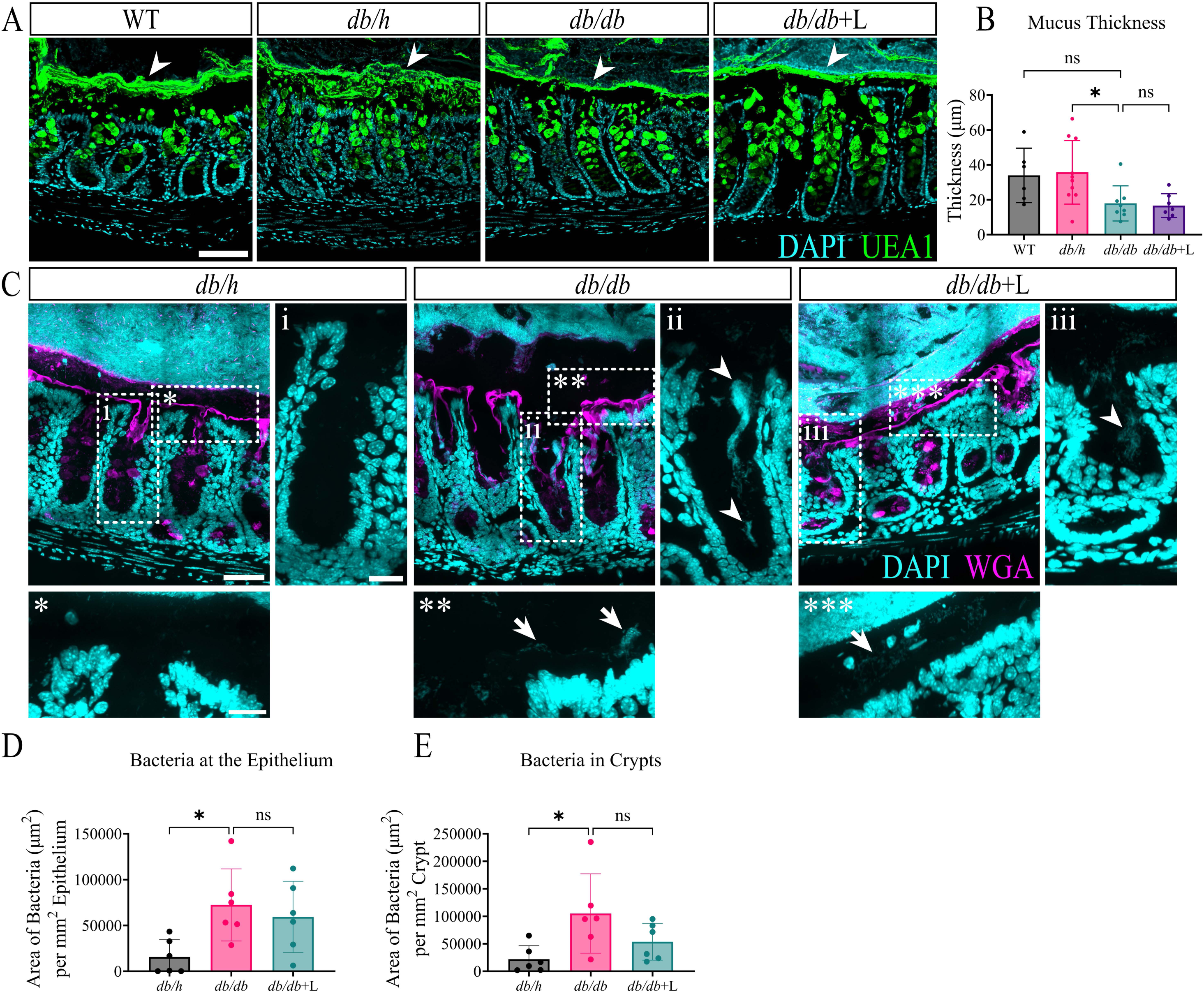
Colonic mucus is thinner and bacteria localize closer to the epithelium in db/db mice. (A) Representative images of mouse colon cryosections labeled for DAPI (cyan) and UEA1 (green) to depict the thickness of the mucus layer between WT, *db/h*, *db/db*, and *db/db*+liraglutide-treated mice. White arrowheads indicate the mucus layer identified by UEA1^+^ labeling. (B) The average thickness of mucus at the epithelium was reduced in the *db/db* colon. Each data point represents the mean of triplicate measurements from an individual mouse. (C) High-resolution images of bacteria located in and around the mucosa detected by labeling with DAPI (cyan). WGA (magenta) labeling was used to define the epithelium. Zoomed in insets of the crypts and epithelium are depicted by i, ii, iii and *, **, *** respectively. White arrows and arrowheads depict bacteria at the epithelium and biofilm formations observed in crypts, respectively. Graphs summarizing the area of bacteria located at the (D) epithelium or in (E) crypts. Data are presented as the mean ± SD, with each data point representing a measurement from an individual mouse. Data were analyzed using a one-way ANOVA with Dunnett’s multiple comparisons test. Scale bars: (A) 75µm, (C) 60µm, 30µm inset. (B) n=6-10 mice per group. (D, E) n=6 mice per group. ns=not significant; *p<0.05.

### Innervation density is reduced throughout the *db/db* colon

Peripheral neuropathy is a known complication in diabetes^2,41^. Neural signaling is an important regulator of intestinal barrier homeostasis, including mucus secretion. To investigate the impact of diabetes on innervation, the colon was immunolabelled with the pan-neuronal marker PGP9.5, and CGRP and SP to distinguish sensory nerve fibers and enteric afferent innervation **(Fig. 4A)**. Total innervation was significantly reduced in the mucosa and muscle layers of the *db/db* colon when compared to WT and *db/h* controls **(Fig. 4B, F)**. CGRP^+^ and SP^+^ nerve fibers were both decreased in mucosa and muscle of the diabetic colon compared to non-diabetic controls **(Fig. 4C, D, G, H)**. CGRP^+^/SP^+^ nerve fibers were also decreased in the colonic mucosa and muscle compared to controls **(Fig. 4E, I)**. Liraglutide treatment had no effect on the *db/db*-associated changes in innervation. These findings demonstrate that colonic innervation is significantly reduced in the *db/db* model of T2DM, with implications for the neural control of intestinal barrier function and mucus release.

**Figure 4:**
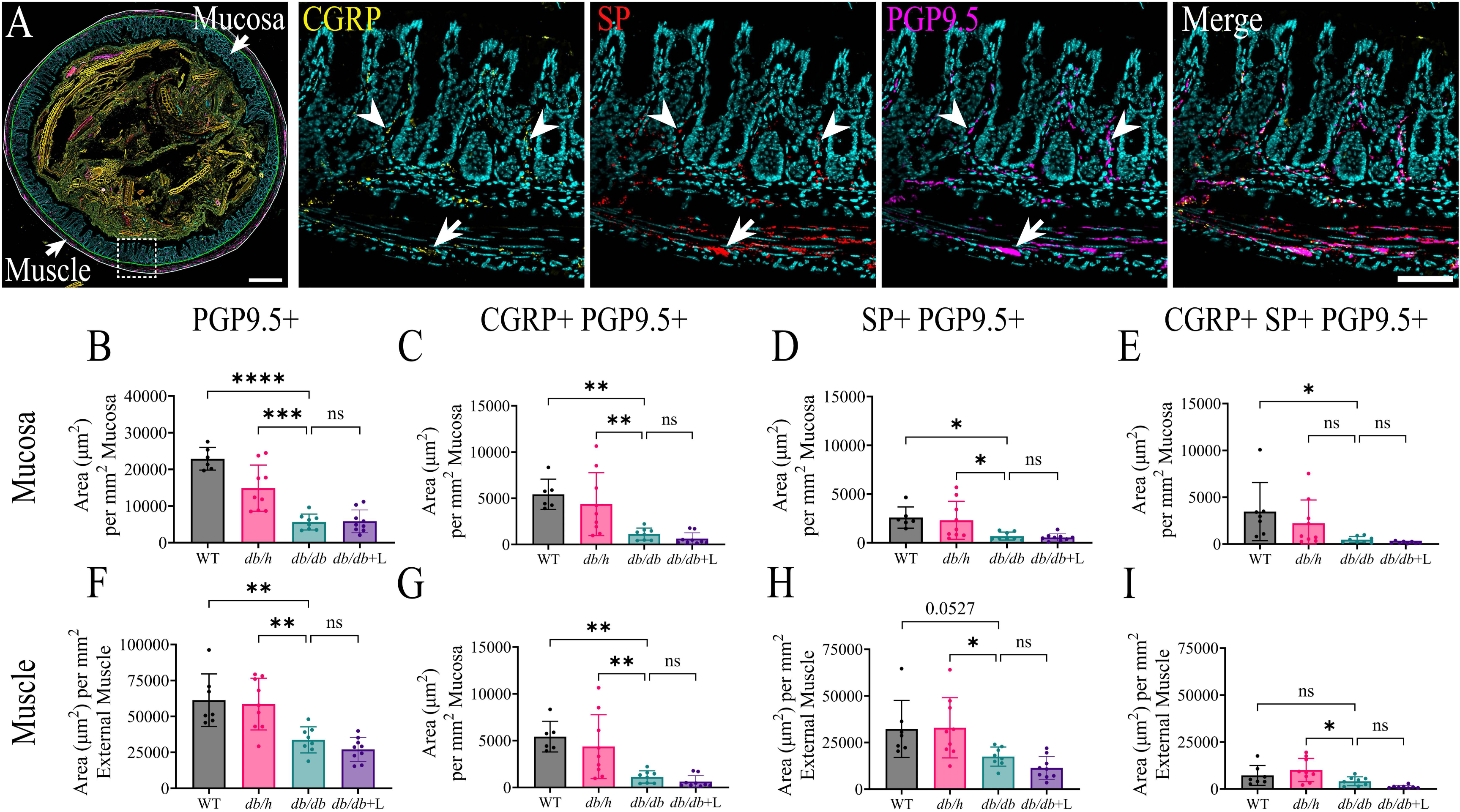
Innervation density is reduced in the db/db colon. (A) Representative image of a *db/h* mouse colon labelled for DAPI (cyan), CGRP (yellow), SP (red) and PGP9.5 (magenta). White arrowheads and arrows indicate innervation in the mucosa and myenteric plexus respectively. Area of PGP9.5^+^ expression in the colonic (B) mucosa and (C) muscle layers of WT, *db/h*, *db/db*, and *db/db*+liraglutide-treated mice. Area of CGRP^+^ PGP9.5^+^ expression in the (D) mucosa and (E) muscle layers. Area of SP^+^ PGP9.5^+^ expression in the (F) mucosa and (G) muscle layers. Area of CGRP^+^ SP^+^ PGP9.5^+^ expression in the (H) mucosa and (I) muscle layers. Data are represented as the mean ± SD, with each data point representing the mean of triplicate measurements from an individual mouse. Data were analyzed using a one-way ANOVA with Dunnett’s multiple comparisons test. Scale bars: (A) 300µm (whole colon), 60µm (inset). n=6-10 mice per group. ns=not significant; *p<0.05; **p<0.01; ****p<0.0001.

### The thickness of the mucosa is increased in the *db/db* colon

Diabetes is associated with chronic low-grade inflammation, which may impact colonic morphology and crypt architecture. Colon tissues were immunolabeled and analyzed to assess mucosal morphology, epithelial cell proliferation, and local inflammation. The mucosa was significantly thicker in the colon from *db/db* mice compared to WT and *db/h* controls **(Fig. 5A, B)**. Treatment of *db/db* mice with liraglutide also reduced the mucosal thickness at the 90% confidence level, when compared to *db/h* control. The diameter of the colonic mucosa was unchanged in diabetes **(Fig. 5C)**. Colon diameter measurements are influenced by undulations in the tissue section whereas mucosal thickness is likely to provide a more reliable measurement of mucosal hypertrophy. The average number of DAPI^+^ cells within the mucosa was unchanged in diabetes. **(Fig. 5D)**. Liraglutide treatment significantly reduced the number of mucosal nuclei. Colon sections were immunolabeled for Ki67 to assess the relative number of proliferative cells in different tissue layers **(Fig. 5E)**. There was no difference in the percentage of Ki67^+^ cells in the mucosa between the *db/h* and *db/db* colon **(Fig. 5F)**. The percentage of Ki67^+^ proliferative cells in the muscle was decreased in diabetes **(Fig. 5G)**. Immunolabeling for 3-nitrotyrosine (3-NT) was used to assess levels of oxidative stress and inflammation in the tissue **(Fig. 5H)**^42^. There was no difference in 3-NT expression in the mucosa or muscle in diabetic mice **(Fig. 5I, J)**. These observations are consistent with hypertrophy of the *db/db* colonic mucosa in the absence of marked inflammation.

**Figure 5:**
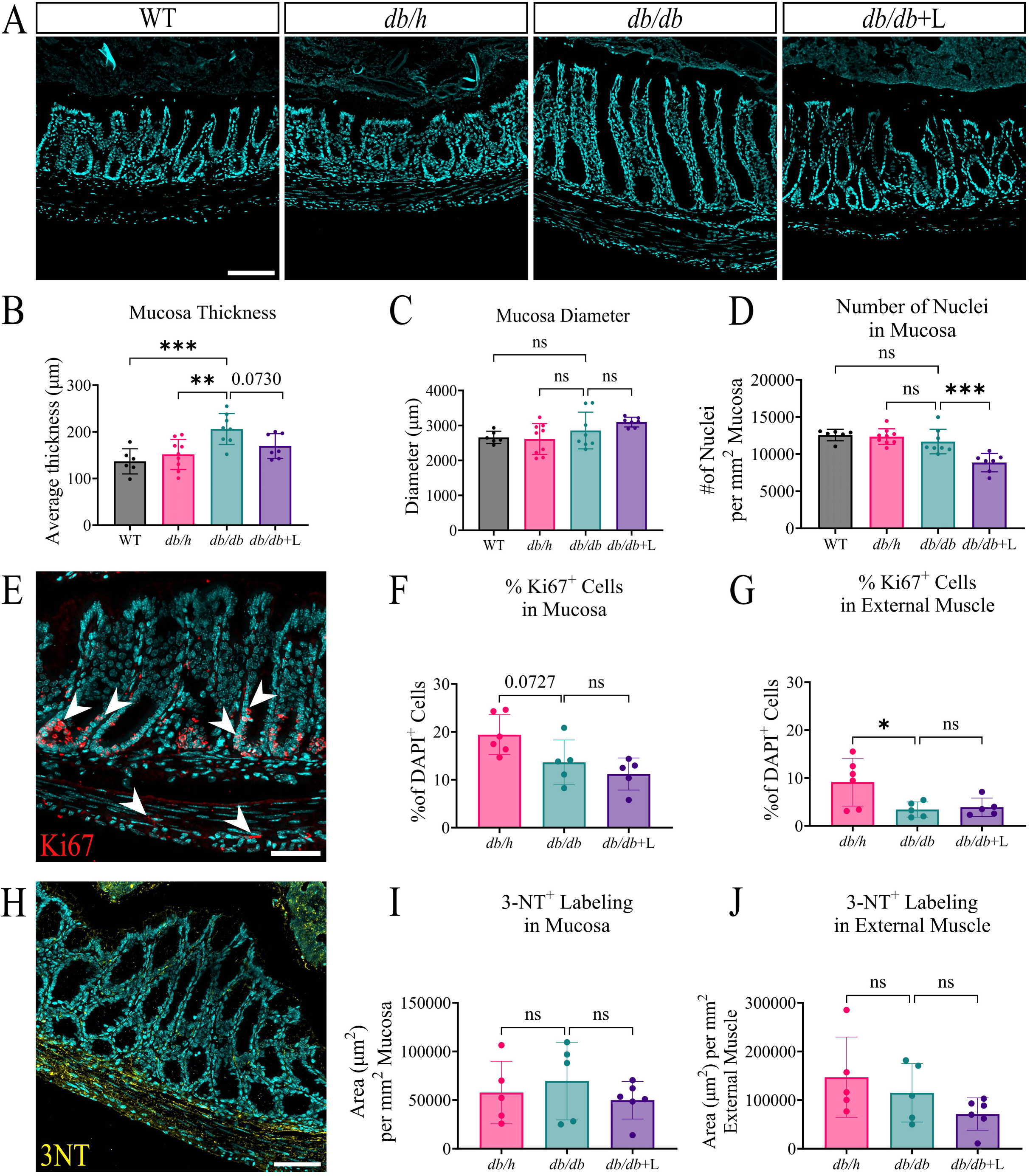
The db/db colonic mucosa is significantly thicker compared to db/h control. (A) Representative images of the WT, *db/h*, *db/db*, and *db/db*+liraglutide-treated mouse colon labeled for DAPI (cyan). (B) Average thickness of the mucosa. (C) Average diameter of the mucosa. (D) Number of DAPI^+^ nuclei in the mucosa. (E) Example image of a *db/h* colon labeled for Ki67 (red) and DAPI (cyan). Percentage of Ki67^+^ nuclei in the (F) mucosa and (G) muscle. (H) Example image of a *db/h* colon labeled for 3-NT (yellow) and DAPI (cyan). Area of 3-NT^+^ expression in the (I) mucosa and (J) muscle. Data are presented as the mean ± SD, with each data point representing the mean of triplicate measurements from an individual mouse. Data were analyzed using a one-way ANOVA with Dunnett’s multiple comparisons test. Scale bars: (A) 100µm, (E) 50µm, (H) 75µm. (B, C, D) n=6-10 mice per group. (F, G, I, J) n=5-6 per group. ns=not significant; *p<0.05; **p<0.01; ***p<0.001.

## Discussion

In this study, we demonstrated an increase in the number of GCs per crypt and a thinner mucus layer in the colon of diabetic *db/db* mice. These changes corresponded to a mucus barrier that was less effective at excluding bacteria from the underlying epithelium. These findings may be explained by disrupted mucus secretion or synthesis. T2DM-associated changes to GC numbers, mucus thickness and bacteria translocation were unaffected by treatment with the clinically used GLP-1RA liraglutide. Accumulation of mucus in GCs or an increase in their overall size was not observed in the *db/db* colon. However, treatment of *db/db* mice with liraglutide was associated with a significant increase in the size of GCs. This may indicate that GLP-1R activation promoted mucus synthesis, but T2DM-associated impaired secretion mechanisms prevented re-establishment of the mucus layer. GLP-1RAs have been reported to influence mucus secretion in a location-dependent manner. The effect of liraglutide to regulate the mucus layer in healthy mice was not assessed in this study due to animal welfare concerns associated with GLP-1RA treatment. Treatment of healthy mice with the GLP-1RA Exendin 4 has been shown to increase mucus secretion in the duodenum, whereas liraglutide inhibits mucus secretion in the lung^43,44^. Although we did not directly assess the functional capacity of GCs to release mucus, previous studies in rats using the streptozotocin model of T1DM have proposed that mucus secretion from GCs is reduced in the small intestine^17^. Defining the impact of T2DM on mucus secretion mechanisms is important to understand how GCs can be targeted to enhance mucus barrier function in patients.

DM patients have a significantly higher risk of infection, including sepsis and infections originating in the gastrointestinal tract^45^. Bacterial contact with the intestinal epithelium is one of the initial steps leading to infection and subsequent inflammation^46^. Impairment or absence of the mucus layer enables bacteria to more readily contact the epithelium and is correlated with ulcerative colitis^34,37,46^. We observed an increase in bacteria translocation to the epithelium and to crypts in the colon of *db/db* mice. Previous studies that have genetically disrupted the mucus layer using *Fut2*^-/-^ or *Muc2*^-/-^ mice demonstrated that bacteria are in close proximity to the intestinal epithelium in the absence of functional mucus^13,47^. Furthermore, this translocation was associated with the formation of microbial biofilms within crypts, consistent with our own observations. These reports suggest that the colonic mucus layer is likely to be an ineffective barrier to bacterial passage in *db/db* mice. The number of intercrypt GCs at the epithelium was significantly decreased in *db/db* mice. Dysfunctional intercrypt GCs have been correlated with a defective mucus layer and increased mucus penetrability using mice lacking SPDEF, a transcription factor required for the terminal differentiation of GCs^34^. Additionally, the number of intercrypt GCs is reduced in the colon of ulcerative colitis patients^34^. Together, this may indicate that alterations to intercrypt GCs may contribute to the diabetes-associated impairment of the mucus layer observed in *db/db* mice. Although we did not observe any significant differences to MUC2 expression within the crypts from the *db/db* mouse colon, it is possible that the expression of other key mucus-associated proteins or peptides integral to mucus integrity, permeability and innate immune responses, such as anterior gradient protein 2, calcium-activated chloride channel regulator 1, or trefoil factor 3 may be reduced. Increased aggregation of bacteria at the intestinal epithelium from *db/db* mice is consistent with clinical observations in DM. Small intestinal bacterial overgrowth (SIBO) is observed in approximately 30% of DM patients and can be detected by a lactulose breath test^48,49^. An impaired mucus barrier may contribute to the high prevalence of SIBO and elevated risk of infection for DM patients. SIBO has also been reported in the streptozotocin rat model of T1DM, and insulin treatment was shown to recover overgrowth to the level of non-diabetic controls^50^. In this instance, insulin dosing commenced immediately after diabetic status was confirmed and may suggest that bacterial infiltration and colonization is not reversible in our T2DM model using the current late stage liraglutide treatment regimen.

Reports of DM-associated alterations to the number of GCs vary between studies^19–21^. We observed a significant increase in the total number of GCs when measured per crypt, but not when measured relative to the whole mucosa. We have also demonstrated that different approaches to GC quantification can lead to different results and thus the specific analysis method used should be considered and clearly described when assessing GC populations. The distribution of GC subsets was shifted such that UEA1^+^ GCs and WGA^+^ GCs were localized slightly closer towards the bottom of the crypt in *db/db* mice, although there was still clear labelling of UEA1^+^ GCs at the apical surface and WGA^+^ GCs at base of crypts. This is consistent with previous evidence that UEA1 and WGA label apical and basal GCs respectively^34^. Together, this may suggest that GC maturation and development from the base through apical surface is unchanged in the diabetic *db/db* colon. We also found that the average length of colonic crypts was significantly increased in the *db/db* mouse and correlated with the increased number of GCs per crypt.

DM is associated with gastrointestinal inflammation^51^. We observed a significant increase in mucosal thickness along with crypt length in the *db/db* colon which may indicate hypertrophy. Increased mucosal thickening has been reported in the small intestine of animal models of T1DM and T2DM, as well as in gastrointestinal tissue from patients with inflammatory bowel disease^12,19,52–54^. However, there were no differences to 3-NT labeling, which detects S-nitrosylated proteins and can indicate tissue inflammation. Elevated S-nitrosylation is associated with insulin resistance and oxidative stress in DM^55,56^. We also observed no difference in the percentage of Ki67^+^ cells in the mucosa which suggests that there is no change to epithelial cell proliferation. In the streptozotocin rat model of T1DM, epithelial cell proliferation has been reported to either increase or not change in the DM small intestine, as determined by BrdU labeling or ornithine decarboxylase activity respectively^53,54^. DM-associated small intestinal mucosa thickening was proposed to be caused by a deficit to apoptotic mechanisms, leading to hypertrophy^54^. Mucosal epithelial cell apoptosis has been assessed in the streptozotocin model of T1DM, but the findings were inconsistent, with reports of increased, decreased, or no change to apoptosis in the T1DM rat jejunum and ileum^53,54^. Alterations to apoptosis mechanisms may explain the increase in crypt length with no change to proliferative cell numbers and thus further investigation is warranted.

Peripheral neuropathy is a major complication of DM and impacts nerves innervating the gut^2^. We observed a reduction in total innervation in the colon of *db/db* mice relative to healthy controls. Additionally, there was a reduction in CGRP^+^ and SP^+^ nerve fibers in the mucosa and external muscle. CGRP is an important regulator of mucus secretion in the colon and depletion of sensory innervation leads to a significantly thinner mucus layer^57^. Previous studies investigating peripheral neuropathy in the *db/db* mouse model have identified behavioral deficits to mechanical and thermal sensation^58,59^. Enteric innervation is also reduced in the colon of DM patients^2^. Our observations suggest a partial loss of sensory nerve fibers and/or enteric afferent innervation in the *db/db* colon. Liraglutide treatment had no effect on the decreased intestinal innervation. GLP-1RAs have been previously reported to have neuroprotective effects, primarily in the context of the central nervous system^60^. However, GLP-1RA treatment has also been shown to increase enteric neuron cell survival *in vitro*^61^. GC functions and maintenance are likely to be regulated by nerves, therefore a reduction to intestinal innervation may contribute to the thinning of the mucus layer^15^. Peripheral neuropathy is also associated with worsening of SIBO in T1DM patients^25^. From our studies we cannot conclude whether a deficit in neuronal input exacerbates or promotes the alterations to GCs and the mucus layer. Although this is beyond the scope of the present study, specific assessment of the mechanistic link between nerves and GC function in DM would greatly enhance our understanding of how the mucus barrier is regulated and potential avenues to preserve or restore function of the barrier in disease.

The effect of GLP-1RA therapies on GCs and the mucus layer during the treatment of diabetes is poorly defined. Our study in a T2DM model of uncontrolled hyperglycemia has provided insight into GLP-1RA-dependent effects on GC size, and suggests negligible effects to GC numbers, mucus thickness and innervation. Understanding the peripheral effects of GLP-1RAs will be valuable in applying these therapeutics for DM-associated conditions beyond glycemic regulation. Furthermore, understanding the effect of GLP-1RAs to influence GCs and mucus in the healthy colon will be essential given the diverse applications of GLP-1RAs. This was not specifically investigated in this study due to animal welfare concerns related to an excessive loss of body weight and lean muscle mass^62^.

## Conclusion

This study has demonstrated that the mucus barrier is impaired in a mouse model of T2DM. We observed increased bacterial colonization at the epithelium and formation of microbial biofilms in the colonic crypts, consistent with an impaired mucus barrier and a higher susceptibility to infection. The increased potential for bacterial translocation across the epithelium aligns with previous reports of an elevated infection risk in DM. Furthermore, both the reduction of intercrypt GCs and peripheral neuropathy may be contributing factors to the mucus layer deficit observed in T2DM.

Treatment with liraglutide effectively managed glycemia but was ineffective in reversing diabetes-associated changes to mucus layer thickness and barrier function. Innervation of the mucosa was not restored following six weeks treatment with liraglutide. Further exploration to understand the impact of GLP-1RA treatment to regulate and minimize T2DM-associated defects to intestinal barrier integrity and innervation may inform the management of T2DM. Maintenance of GCs and the mucus layer is critical for an effective intestinal barrier to minimize the risk of infection in diabetes.

## Methods

### Animal studies

Mice with T2DM induced by a deficiency in the leptin receptor (*db/db*) and their heterozygote littermates (*db/h*), were bred by the Monash Animal Research Platform, using Jackson Laboratory mice (strain #000697: B6.BKS(D)-Lepr^db^/J) (10-week-old, male and female). *Dock7^m^* +/+ mice were used for wild-type (WT) controls (strain #000642: BKS.Cg-Dock7m +/+ Leprdb/J). All mice were housed under a 12 h light/dark cycle, controlled temperature (24°C), with free access to food and water. All procedures involving mice were approved by the Monash Institute of Pharmaceutical Sciences animal ethics committee (#39545). Studies involving animal experiments conformed to the Animal Research: Reporting of In Vivo Experiments (ARRIVE) guidelines. From 16-weeks of age, *db/db* mice were treated with liraglutide (1mg/kg) or saline via subcutaneous injection three times per week. Liraglutide treatment controls were not performed on *db/h* mice due to welfare concerns of excessive loss of weight and lean muscle mass. For in vivo experiments, mice were randomly assigned to treatment groups, investigators were blinded, and equal group sizes were obtained to minimize bias. Diabetic status was confirmed by blood glucose measurements once every two weeks, using a handheld glucometer (Accu-Chek® Performa; Roche Diagnostics) where the upper limit of detection for the glucometer was 33.3 mmol/L; any readings above this value were recorded as 33.3 mmol/L. Mice with blood glucose measurement above 13.3mmol/L were considered diabetic. At study endpoint the distal colon was collected.

### Tissue Preparation

Distal colon tissue was fixed in 4% paraformaldehyde (PFA) at 4°C for 24 h. PFA was then cleared by washing with phosphate buffered saline (PBS) and the tissue was cryoprotected in 30% sucrose (w/v in PBS) for a minimum of 24 h before snap freezing in OCT compound and cryostat sectioning (10µm thickness) on a Leica CM1950 cryostat. Sections were cut from colon regions where a fecal pellet was intact. The presence of the fecal pellet is important for stabilizing the mucus layer, allowing for mucus thickness to be assessed by immunostaining.

### Immunostaining

Cryosections were air dried on Superfrost Plus slides and incubated in blocking buffer (5% normal donkey serum, in PBS containing 0.1% Triton X-100 and 0.1% sodium azide) for 30 min to permeabilize the tissue and minimize nonspecific binding of antibodies. Sections were then incubated with primary antibodies **(Table 1)** (overnight at RT in blocking buffer). Excess primary antibodies were washed off with PBS (3 x 10 min) and sections were then incubated for 1 h at RT with secondary antibodies prepared in PBS **(Table 2)**. Unbound secondary antibodies were washed off with PBS (2 x 10 min), then sections were incubated with the nuclear marker DAPI (1: 1000 in PBS, 5 min at RT; Sigma Aldrich). Tissues were washed (1 x 10 min PBS), then incubated with Vector TrueVIEW Autofluorescence Quenching solution (Vector Laboratories) for 5 min at RT. Sections were then washed (1 x 10 min PBS) before mounting with ProLong Diamond Antifade mounting medium (Thermo Fisher Scientific).

**Table 1:**
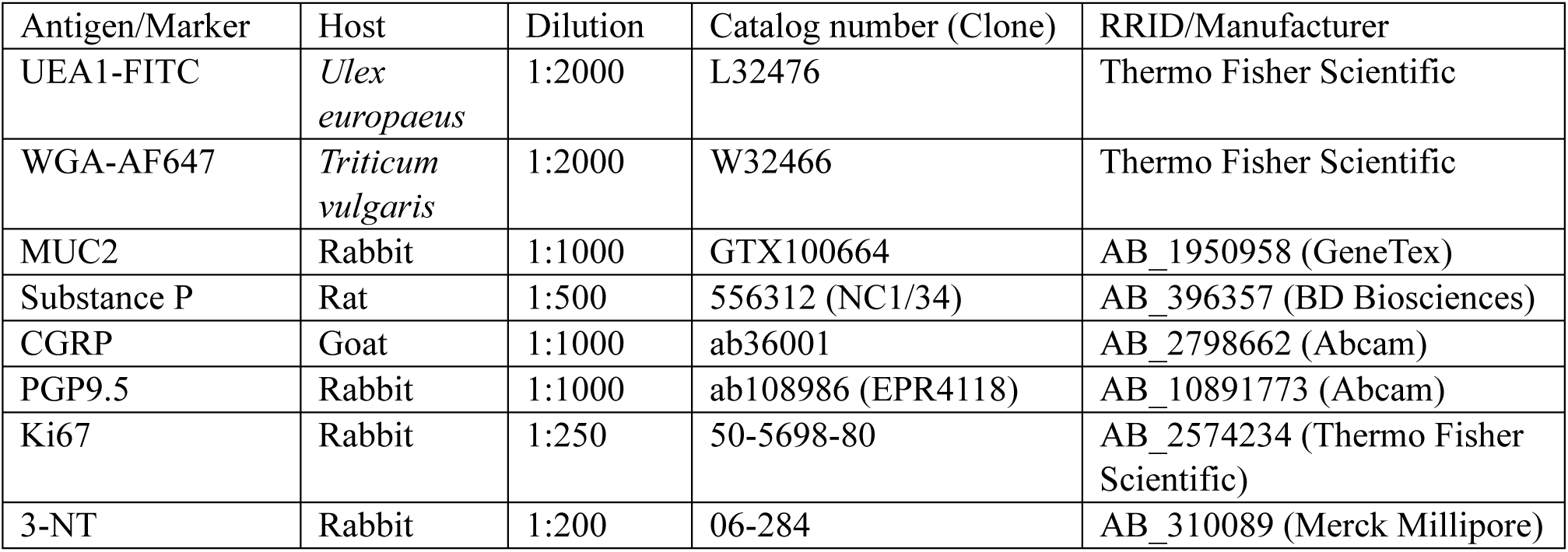
Primary antibodies used in this study.

| Antigen/Marker | Host | Dilution | Catalog number (Clone) | RRID/Manufacturer |
| --- | --- | --- | --- | --- |
| UEA1-FITC | *Ulex europaeus* | 1:2000 | L32476 | Thermo Fisher Scientific |
| WGA-AF647 | *Triticum vulgaris* | 1:2000 | W32466 | Thermo Fisher Scientific |
| MUC2 | Rabbit | 1:1000 | GTX100664 | AB_1950958 (GeneTex) |
| Substance P | Rat | 1:500 | 556312 (NC1/34) | AB_396357 (BD Biosciences) |
| CGRP | Goat | 1:1000 | ab36001 | AB_2798662 (Abcam) |
| PGP9.5 | Rabbit | 1:1000 | ab108986 (EPR4118) | AB_10891773 (Abcam) |
| Ki67 | Rabbit | 1:250 | 50-5698-80 | AB_2574234 (Thermo Fisher Scientific) |
| 3-NT | Rabbit | 1:200 | 06-284 | AB_310089 (Merck Millipore) |

**Table 2:** Secondary antibodies used in this study.

| Target | Host | Dilution | RRID |
| --- | --- | --- | --- |
| α-rabbit AF568 | Donkey | 1:500 | AB_2534017 |
| α-rabbit AF750 | Donkey | 1:250 | AB_2943056 |
| α-rat AF647 | Donkey | 1:500 | AB_2910635 |
| α-sheep AF568 | Donkey | 1:500 | AB_10055702 |

### Image Acquisition and Microscopy

Images were captured on a Leica DMi8 widefield microscope. Tile scan images with Z-steps (15.93µm, 0.693µm steps) were acquired using a 20x air objective (0.8NA). The resolution of each tile was 2048x2048 pixels, with a pixel size of 0.325µm^2^. Acquired widefield images were deconvolved using Huygen’s professional deconvolution software (version 25.04.0p1) prior to analysis.

Images of bacterial clusters were acquired on a VT-iSIM super-resolution microscope (VisiTech International). Tile scanned images with Z-steps covering the full thickness of the tissue were acquired using a 60x silicon oil objective (1.30NA). The resolution of each tile was 1469x986 pixels, with a pixel size of 0.108µm^2^.

### Image Analysis

Fiji open-source software (version 1.54f) was used for quantitative analysis of immunofluorescent images^63^. Custom Fiji macros were written to automate the image analysis **(Fig. S2)**^64^. Briefly, individual GCs and positively labeled area of each fluorescent marker was segmented used custom-trained ilastik and Cellpose models^65,66^. Segmented GCs smaller than 10µm^2^ were excluded from the analysis. The area of the whole mucosa or external muscle was also measured and then used to normalize results per mm^2^. For crypt-specific analysis, the analysis was restricted to manually defined regions of intact crypts that span from the base of the crypt to epithelium **(Fig. S4)**. A minimum of seven intact crypts was measured per mouse. For bacterial cluster analysis, manual thresholds were used to segment DAPI^+^ bacteria area due to variability in image intensities. The area of bacterial clusters was then normalized to the area of the epithelium and crypts to calculate bacteria area per mm^2^.

Ilastik open-source software was used to segment the positive labelling from each channel into binary masks^65^. Specific pixel classifier models were trained to identify staining of mature MUC2, UEA1, Ki67, 3-NT, CGRP, SP and PGP9.5 in tissue sections. Ilastik-segmented binary masks were used to measure the area of positive staining and mucus layer thickness. Twelve to twenty-eight deconvolved maximum projection stacks sampling control, vehicle and treatment group mouse tissues, were used to train each machine learning model. Images were manually annotated to define positive staining and background signal, and the training was validated on 5 images that were not used for the training. The trained pixel classifiers were used as segmentation models in the analysis.

Cellpose was used to segment and count individual GCs^66^. Cellpose is a deep learning neural-net based algorithm used for cell segmentation. A Cellpose model was trained to detect UEA1^+^ and WGA^+^ GC labelling in the deconvolved images of mouse colon. Fifty-eight images were included to train the GC model using a human-in-the-loop approach. The ‘cyto’ pretrained model was used to approximate segmentation and then images were manually annotated to refine the selection of individual GCs. The segmentation model were validated on five images that were not used for the training.

### Statistical Analysis

Statistical analysis was performed using GraphPad Prism (version 10.6.1). Data was analyzed using a one-way ANOVA with Dunnett’s multiple comparisons test with *db/db* data as the comparator. A p value below 0.05 was considered significant. The specific statistical tests used, and numbers of replicates are also provided in the respective figure legends.

## Abbreviations

3-NT: 3-nitroyrosine
CGRP: Calcitonin Gene-Related Peptide
DM: Diabetes Mellitus
GCs: Goblet cells
GLP-1: Glucagon-like Peptide 1
GLP-1R: Glucagon-like Peptide Receptor
GLP-1RA: Glucagon-like Peptide 1 Receptor Agonist
MUC2: Mucin-2
PFA: Paraformaldehyde
PGP9.5: Protein Gene Product 9.5
SIBO: Small Intestinal Bacterial Overgrowth
SP: Substance P
T1DM: Type 1 Diabetes
T2DM: Type 2 Diabetes
UEA1: *Ulex europaeus* agglutinin 1
WGA: Wheat germ agglutinin)

## Acknowledgments

The authors thank Alana Risteska for her contribution to the image analysis of diabetic tissue sections.

---

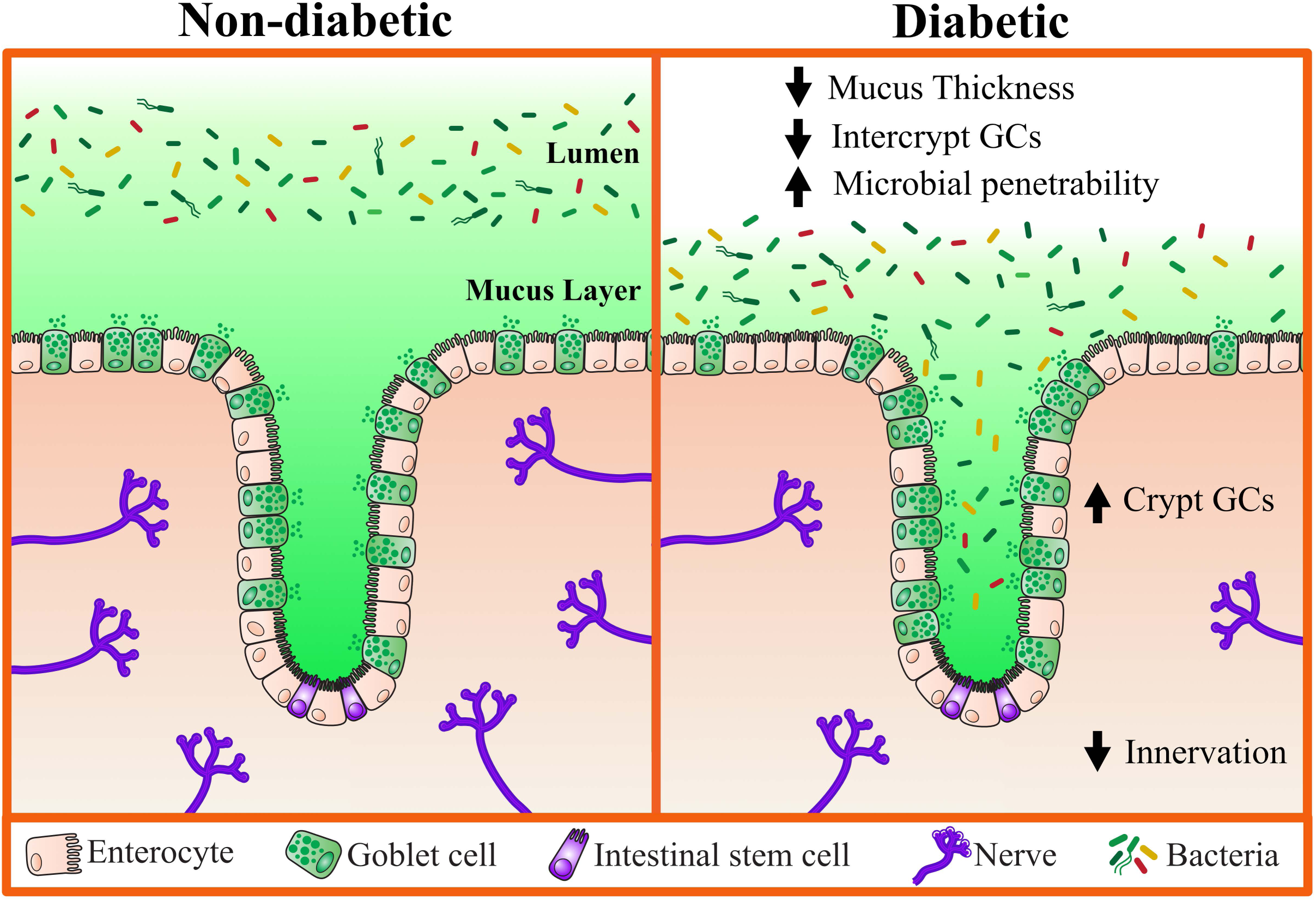

