## Supplementary Data for "The Colonic Mucus Layer is Thinner and is Associated with Goblet Cell Hyperplasia in the *db/db* Mouse Model of Type 2 Diabetes"

Rowe MC *et al.*

**Supplementary Information**

**Analysis workflow to define GCs and mucus.**

Automated and unbiased image analysis is critical for large scale quantification of microscopy images. A custom image analysis workflow was developed in Fiji to quantify colonic GC numbers and mucus thickness from captured images. Deconvolved widefield images were manually annotated to define key regions for analysis including the mucosa, mucus layer, and individual crypts **(Fig. S2A, B)**. GCs and the mucus layer were segmented using custom-trained Cellpose and Ilastik models respectively **(Fig. S2C, S3)**. The number, size and area of each GC marker was calculated from the segmented GC masks. Additionally, GCs located at the epithelium were identified as intercrypt GCs. Analysis was subsequently restricted to intact individual crypts that span from the base of mucosa to the epithelium, and the same measurements were calculated **(Fig. S2D, E, S4)**. The location of individual GCs within each crypt was also measured **(Fig. S2F)**. The workflow described streamlines image analysis of intestinal tissue sections.

**Supplementary Figure Legends:**

**Fig S1:** ***Experimental timeline.*** WT, *db/h* or *db/db* mice (10-wk-old, male and female) were maintained on a normal chow diet. Blood glucose and body weight were measured every two weeks throughout the experimental timeline. From 16 weeks of age *db/db* mice received liraglutide injections (1mg/kg, s.c., 3x weekly) or vehicle control for 6 weeks and were then humanely killed at endpoint for distal colon collection. ****$p<0.001$.

**Figure S2:** ***Goblet cell image analysis workflow.*** (A) Deconvolved widefield images of mouse colon cross sections labeled with DAPI (cyan), UEA1 (green), MUC2 (yellow), and WGA (magenta). (B) Manual annotation to define key regions of interest including the mucosa, mucus layer, and individual crypts. (C) An example inset of the annotated image and subsequent segmentation of $UEA1^+$, $WGA^+$ and $MUC2^+$ labeling used to count the number of GCs and measure the thickness of the mucus layer. (D) Example of an individual crypt selected for crypt-specific analysis. (E) Individual markers and subsequent segmentation masks of UEA1, WGA and MUC2. (F) GC location is measured relative to the base and opening of the crypt. Scale bars: 300µm (whole colon), 75µm (inset), and 25µm (crypt).

**Figure S3:** ***Validation of goblet cell segmentation using Cellpose.*** Example images of UEA1 and WGA labeling in the mouse colonic mucosa. Unique colors represent individual GCs segmented using the custom-trained Cellpose model. Five unique samples across WT, *db/h*, *db/db*, and *db/db*+liraglutide-treated mice that were used to validate the model by comparing the number of GCs segmented manually and with the Cellpose model. Scale bars: 25µm.

**Figure S4:** ***Manual annotation of intact and non-intact crypts.*** Example images of crypts in the mouse colonic mucosa labeled against DAPI (cyan), UEA1 (green) and WGA (magenta). (A) Intact crypts span the full thickness of the mucosa and were selected for crypt-specific analysis. (B) Non-intact crypts do not span from the base of the mucosa to the epithelium and were not selected for crypt-specific analysis. White outlines indicate individual crypts. Matching DAPI-only images are depicted below each image. Scale bar: 50µm.

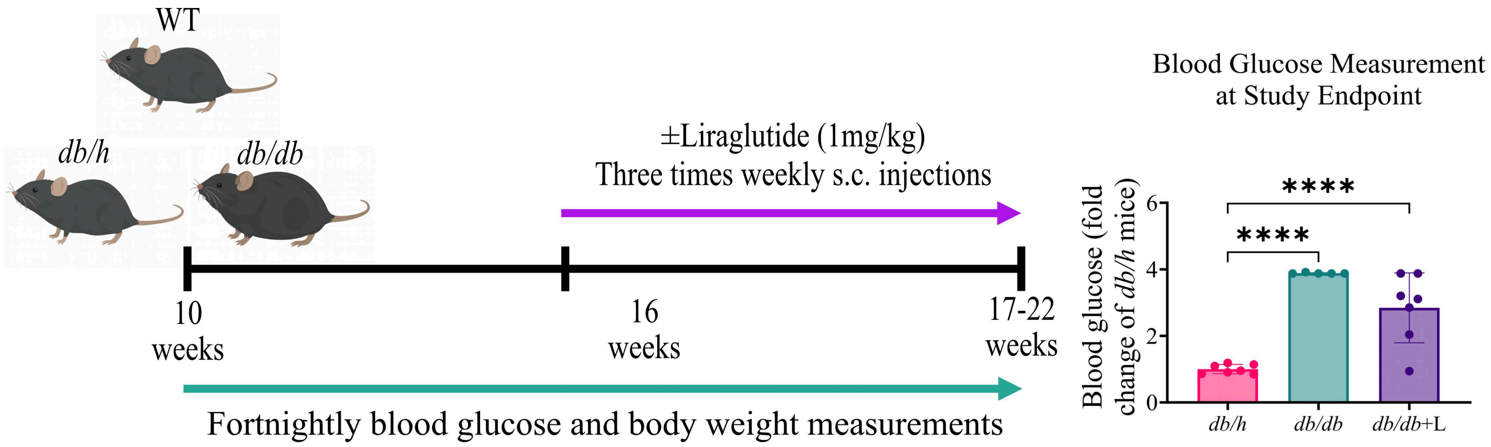
WT
db/h
db/db
±Liraglutide (1mg/kg)
Three times weekly s.c. injections
10
weeks
16
weeks
17-22
weeks
Fortnightly blood glucose and body weight measurements
Blood Glucose Measurement
at Study Endpoint
Blood glucose (fold change of db/h mice)
****
****
6
4
2
0
db/h
db/db
db/db+L

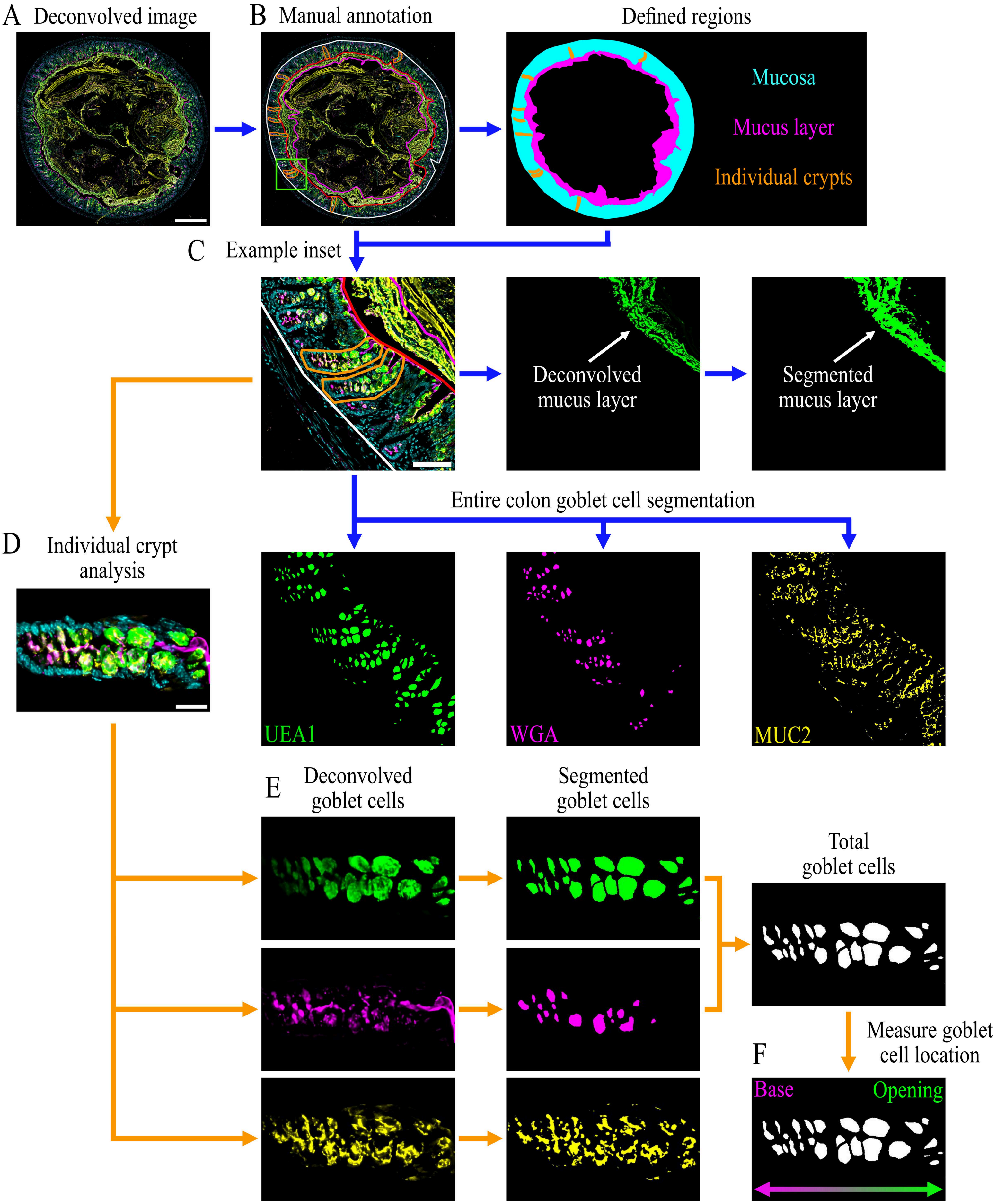

A Deconvolved image
B Manual annotation
Defined regions
Mucosa
Mucus layer
Individual crypts
C Example inset
Deconvolved mucus layer
Segmented mucus layer
Entire colon goblet cell segmentation
D Individual crypt analysis
UEA1
WGA
MUC2
E Deconvolved goblet cells
Segmented goblet cells
Total goblet cells
Measure goblet cell location
F
Base
Opening

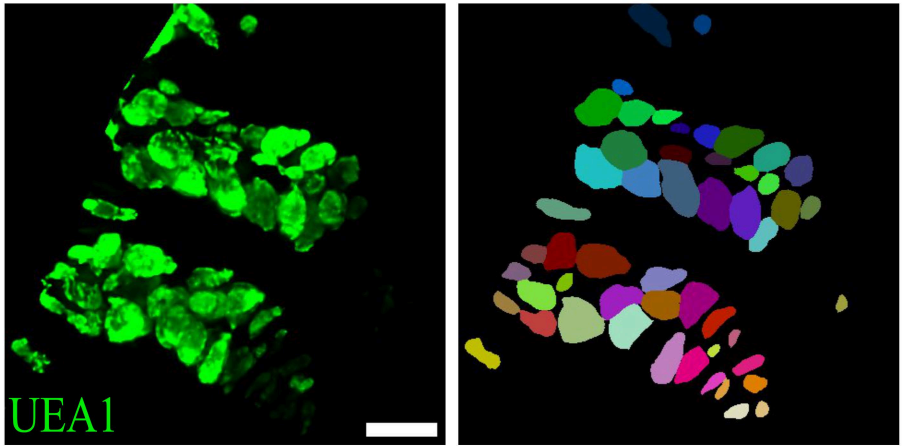


| UEA1 Sample # | # of GCs manually counted | # of GCs counted by Cellpose | % Accuracy |
|---|---|---|---|
| 1 | 55 | 52 | 94.5 |
| 2 | 76 | 78 | 97.4 |
| 3 | 45 | 46 | 97.8 |
| 4 | 81 | 89 | 91.0 |
| 5 | 46 | 43 | 93.5 |

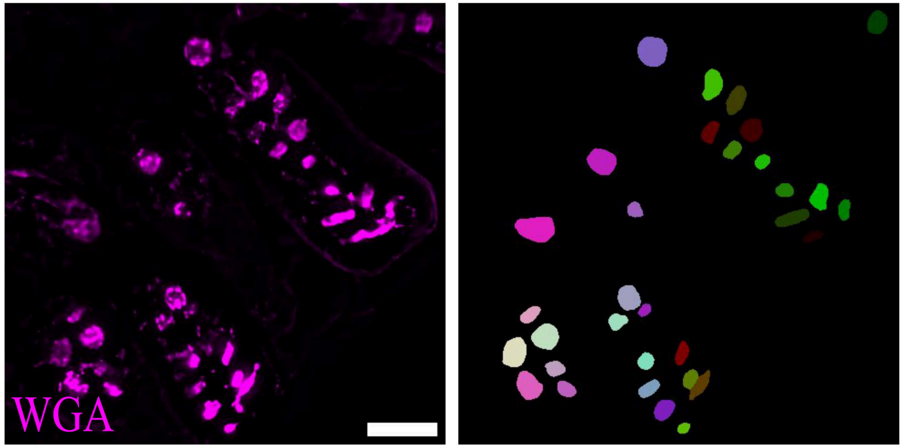


| WGA Sample # | # of GCs manually counted | # of GCs counted by Cellpose | % Accuracy |
|---|---|---|---|
| 1 | 19 | 18 | 94.7 |
| 2 | 32 | 32 | 100 |
| 3 | 22 | 21 | 95.5 |
| 4 | 26 | 25 | 96.2 |
| 5 | 23 | 25 | 92.0 |

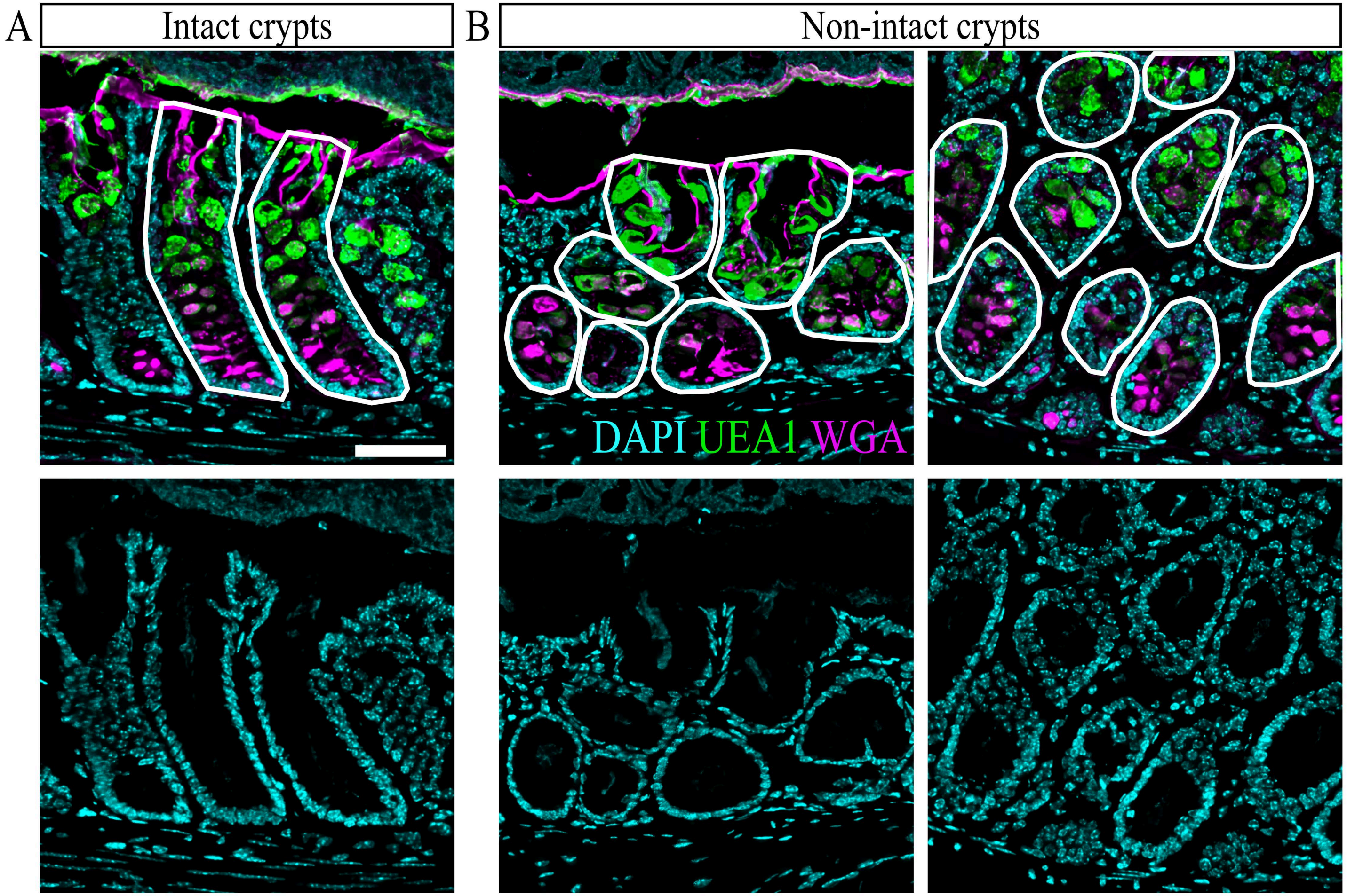
A
Intact crypts
B
Non-intact crypts
DAPI UEA1 WGA
